# The personal space and the collective behavior of crowd disasters

**DOI:** 10.1101/2024.09.05.611443

**Authors:** Igor Lugo, Martha G. Alatriste-Contreras

## Abstract

The personal space in dense crowd situations is commonly underestimated. However, the self-awareness of it can prevent and handle risk situations. The aim of this study was to explore theoretically the use of the personal space as a key concept for designing simple computational models related to collective behaviors and crushing events. We used an agent-based model related to transitional rules associated with the Shelling’s spatial proximity model of segregation. Based on an explorative data analysis and model validation, we found that the dynamics of crowd events showed significant statistical regularities between dense situations and the individual perception of the personal space. These results suggests that crushing events in dense crowd situations are highly probable and that the density levels are the key for delaying the presence of such disasters.

## Introduction

The dynamics of extremely dense crowds or crowd turbulence is a deeply worrying phenomenon in which people have been hurt and lost their lives in different places and cultures. It is characterized by sudden variations in all directions among people that push others around generating uncontrollable and unpredictable large-scale events (Helbing et.al., 2006, 2007). Such events can be triggered by a small group of individuals in close proximity to one another without any intention of hurting people. Unfortunately, the most recent well-known events in Etah, Uttar Pradesh, India, July 2, 2024, during a religious event, or in South Korea on October 29, 2022, during a Halloween celebration, showed a disturbing lack of knowledge required to understand and prevent fatal injuries. In particular, the event in South Korea showed the crush of a crowd without an ex-ante mass panic (Helbing et.al., 2007). Contrasting with these lamentable events, there are other examples in which the overcrowding did not result in tragedy. For example, the celebration of the winning of the soccer world cup by Argentina, at December 20, 2022. In this event, we could see self-organized crowds across the country without fatal crush events. Both incidents were separated by two months apart, and they showed completely different outputs. Therefore, our investigation proposes a theoretical and an interdisciplinary approximation for modeling the dynamics of crowd crushing in social gatherings based on the Shelling’s model of segregation (Schelling 1978; Silver et.al., 2021) . We are particularly interested in studying one specific cultural driver that can explain the individual reaction in dense crowded situations: the personal space (PS). To achieve this, we shall use an agent-based model (ABM) and a group behavior formulation for simulating and showing complex and emergent behaviors in dense crowded situations.

Based on the group behavior formulation Lewin (1945); Yang et.al., (2020), which considers the social cohesion as the key term for understanding interactions between individuals, we use the sociocultural approach and the complex systems perspective to associate a possible cultural driver with the concept of the PS (Vygotsky 1978, 1986; Barabasi 2002; Bar-Yam 2004). This concept is commonly used in studies of human behavior (Gifford 2012; Bonnes and Carrus 2017), and it can be understood as the surrounding area for social interactions of an individual (Welsch et al. 2019; Gifford 2012; Battle 2012). In addition, this PS can be associated with levels of psychological comfort or discomfort (Thompson et al. 1979; Welsch et al. 2019). Consequently, we aim to translate this conceptual idea into the ABM by generating simple local rules on a flat surface without obstacles that can replicate the conditions and dynamics in different venues of dense crowded situations, such as music concerts, religious meetings, and sport events (Helbing et al. 2003). Then, if we consider the PS as a principal issue to understand the presence of restricted physical movements and the collective and individual psychology—which depends on the cultural context, such as family, friends, communities, and religion— of dense crowds, we can describe the instinctive reaction of a person (Reicher 1984). Therefore, we are interested in the following questions: What are the density situations in which the crushing is unstoppable? How probable are the crushing events in such crowded gatherings? What are the perceived levels of comfort or discomfort for being surrounded by similar persons in such situations? We shall answer these questions by using an interdisciplinary approach in science. Just as we mentioned before, we use the sociocultural approach as part of the group behavior formulation for indicating how the daily interpersonal interactions affect our natural instinct for self-preservation in different crowd situations (Battle 2012, 1998; Kelly et al. 2015). That is, social interactions guide and teach individuals how to react instinctively to harmful events. In addition, we use an ABM based on the Shelling’s model of segregation for describing how individuals utilize their perceived PS, interact with each other, and generate large-scale collective behaviors related to crushing events (Schelling 1978). Even though this type of ABM is not commonly used in modeling crowd dynamics, it shows the advantage of generating well-funded and simple models for studying crushing events. Next, we shall use an explorative data analysis and model validation for evaluating our model outputs (Sargent 1984; Kerr and Goethel 2014). In particular, we design an explorative data analysis for clarifying the importance of the PS and the presence of different levels of density situations in the formation of crushing events. Furthermore, the model validation is intended for predicting the existence of crushing events under those initial conditions that are outside of the realm of observed conditions (Lugo and Martínez-Mekler 2022; Lugo and Alatriste-Contreras 2022). Then, we use a probabilistic framework based on a Monte Carlo approximation that looks for statistical regularities associated with the dynamics of such initial conditions.

Therefore, we hypothesize that fatal crushing events can be related to cultural aspects of some regions. Societies that are related to high tolerance thresholds of being surrounded by people in near-body proximities are more vulnerable to fatal crushing in overcrowded events. That is to say, if a person tolerates the violation of the PS, it is more probable that she will be in a fatal crushing. On the other hand, societies that are related to less tolerance thresholds of the PS are less vulnerable to crushing. That is, if a person has less tolerance of the discomfort associated with the violation of the PS, the presence of fatal crushing is less probable. Therefore, the cultural context related to the PS affects the presence of fatal or non-fatal cases of crowd crushing. The practical relevance of this study is to communicate to attendants of dense social gatherings the higher risk of being crushed and to prevent uncontrollable crowd turbulences and their fatal crushing events.

The remaining part of the article proceeds as follows: Literature review, Material, Methods, Results, and Discussion. The Literature review shows the main references about studies of collective behaviors and crowds in different disciplines, as well as their computational approximation. The Material section indicates the programming libraries for coding, generating, and analyzing our ABM. The Method section explains our ABM based on the Shelling’s model of segregation that describes the configuration and transitional rules in different situations related to the PS. Furthermore, we describe the computation of a main measure of nearest neighbors, the implementation of density scenarios, the procedure related to the data analysis and model validation, and the initial conditions. The Result section displays our findings based on the model validation and sensitivity analysis. Finally, the Discussion section shows our final remarks and conclusions.

### Literature review

In the diverse and extended literature related to collective behaviors, we can identify different methods for crowd simulation and modeling. Following the work of Yang et. al., Yang et.al., (2020) and Dang et. al., Dang et. al., (2024), we agree that there are different levels of modeling crowd, for example microscopic and macroscopic levels, and they depend on the scale of analysis. Even though the existence of such approaches that are commonly used separately for describing collective behaviors, they have to be used together for building effective descriptions of the crowd dynamics in question. To be effective, models have not to be necessarily realistic in the sense of describing every detail of real life situations, but instead have to be the tools for “providing insightful explanations of known facts, and revealing the unrealized consequences of widely held beliefs.” (Klosterman 2012). Therefore, we use a simple approximation of modeling in which we use the dynamic group behaviors as the formulation to glue together scales, methods, and ideas of crowd modeling and the PS (Lewin 1945; Dion 2000).

One important characteristic of the group behaviors approach is the type of coordinations and the trigger events that stress them. In particular, studies in the biology field, such as flocking of birds, point out the dynamics behind the effect of local actions and the formation of large-scale patterns within groups (Ioannou & Laskowski 2023). A feature that is highlighted in these studies is the efficient information transfer for preventing crushing events (Sumpter 2006). This is because there is sufficient space between each individual and its near neighbors at the level of the group that communicates effectively the sudden changes of direction and prevents any type of crushing incidents. Even if those changes were generated by warning sings of possible predator around the group, the type of group reaction is to avoid the dangerous situation by impulsive but coordinated dispersion. Therefore, the physical space between individuals in animal groups stands as one key for coordinating movements and prevents crashing or crushing events.

On the contrary, collective behaviors related to groups of people show singular attributes of crowd dynamics (Helbing et.al., 2007; Üsten et al. 2022). In particular, we are interested in the chaotic and uncoordinated movements and their unknown trigger factors. In this respect, there are studies that have considered empirical and theoretical approaches to describe the dynamics in the motions of extremely dense crowds. The work of Helbing et.al., (2007) analyzed video recordings of the crowd disaster in the “Muslim pilgrimage in Mina/Makkah, Saudi Arabia.” They identified turbulent flows that are similar to earthquakes because they initiated as sudden changes of pressure, and these changes generated unstoppable and chaotic movements in the crowd. In addition, the work of Helbing et al. (2003) studies the evacuation dynamics of classrooms by experiments and simulations. They identified jamming of students at the exit door that affected the evacuation dynamics. In this study, the trigger event was a signal alarm shouted by the cameraman. In addition to these findings, the study of Nakayama et al. (2005) analyzed two-dimensional formulation of pedestrian and granular flows. They suggested the presence of instabilities in pedestrian flows. Therefore, in this type of approximation, it is obvious by now that the density level is one other key for understanding crowd disaster.

With respect to the method, the work of Yang et.al., (2020) reviewed closely the models used in crowd simulation. They suggested that the presence of a mesoscopic level of modeling can improve the simulation of more realistic crowds. In the same line, the work of Dang et. al., (2024) extensively reviewed models of pedestrian simulation in high-density situations. They pointed out the importance for considering different levels and integration models for simulating crowds. Compared with these method approaches, the work of Schelling (1978), which we use in this study, about segregation can still offer a simple alternative approach for modeling crowd behaviors. The Shelling’s model of segregation shows the fundamental principles of individual decisions based on their self-interest and group preference (Batty 2005). This model commonly uses the cellular automata (CA) framework for generating ABMs (Batty 2005). Consequently, the Shelling’s model describes crowd behaviors based on a simple psychological process of social groups. In addition, it is worth to mention few studies related to the use of CA in modeling crowd events. For example, the work of Fukui and Ishibashi (1999) suggested the use of CA models for understanding pedestrian flows and “the high density crowd behavior in overcrowding places.” Moreover, the work of Muramatsu and Nagatani (2000) and Burstedde et al. (2001) showed the application of the CA in pedestrian dynamics. They used the dynamics of the traffic flow for describing the pedestrian movements. They identified that the jamming transitions emerge from an increased density. Therefore, these methods indicate the importance to incorporate simple and significant factors for developing models of known facts about collective behaviors.

In line with the analysis of the PS applied to group behaviors, there are very few published studies that point out the need to carefully consider its local effect in different groups of people. For example, Welsch et al. (2019) used a field-theoretical framework for showing the response of the violation of the PS. They found that such a violation is immediate. Then, if your PS constantly changes, there is no tolerance for violations. Compare to this results, the work of Thompson et al. (1979) suggested the existence of a comfort or tolerance zone, around 180–240 cm. Finally, the work of Lugo and Alatriste-Contreras (2022) used the PS concept to validate the application of the classic Moore neighborhood in the CA. They suggested that the PS is a fundamental concept for modeling the interpersonal distance in spatial interactions. Therefore, these studies show the potential to complement the findings of group behaviors and modeling by including explicitly the PS in theoretical and empirical analyses.

Thus far, these references indicate the usefulness of considering an interdisciplinary approach that integrates new ideas into different scales and methods of analysis. In particular, the PS should be part of the literature related to crowd simulation and modeling. It seems that the PS is a fundamental concept for understanding the crushing events. In the section that follows, we present the materials related to programming libraries for using in our ABM.

## Materials

Our ABM is intended for simulating known facts of crushing events in different levels of density scenarios, therefore the materials that we use are related to different programming libraries. The programming code is one of the best tools for exploring the presence of simple local rules, which cannot be seen in real live behaviors, for explaining complex outputs in a crowd. In particular, we apply the Python programming language and its third party libraries for generating our ABM, analyzing its outputs, and showing its results. In particular, we used NumPy, matplotlib, and SciPy libraries. For making all our data, code, and results as open and accesible as possible, we place them in the Open Science Framework (OSF). The name of the project associated with the OSF is Cultural drivers for understanding crowd crushing. Therefore, for a detailed specification of the programming code, see the OSF project.

## Methods

As we mentioned before, we use the Shelling’s model of segregation for generating our ABM (Schelling 1978). Compared with complex models (some of them mentioned in the literature review section) that try to describe reality with all the details, our proposed model shall identify core assumptions and a key variable that can represent the behavior of the system in question. In particular, we assumed that each individual shows a self-awareness of its PS in different scenarios of crowd density and that the computation of a nearest neighbors measure provides the basic information to understand and describe the presence of crushing events in flat areas, indoors or outdoors, without physical obstacles such as chairs, guardrails, threes, stairs, etc. Therefore, we used the Shelling’s model of segregation as a vehicle for tracing through the implications of the effects of the PS related to two types of individuals, tolerant or intolerant of being sounded by similars, in different levels of crowd density.

Our ABM describes individuals who change location or stay in the same place according to the presence of individuals with different or similar levels of tolerance. We define tolerance or intolerance in terms of the PS as the individual perception of comfort or discomfort associated with the number of individuals around with similar perception. Therefore, our model describes a spatial model of segregation in which an individual will change its location as a response of the sense of discomfort associated with the presence of surrounding individuals, or an individual will stay in place as a response of aligning common preferences of surrounding individuals. The following subsections describe in detail the configuration of our model and the individual behavior of being surrounded by others, as well as the proposing data analysis and model validation.

### Configuration

To build our theoretical ABM model, we used different references for defining the configuration of a matrix, the attributes of each cell, and the shape of the neighborhood of an individual. In this study, we use the basis of CAs for modeling two-dimensions of crowd events that can represent a wide variety of dynamic systems related to crowd modeling and its physical and temporal scales (Janelle 2005). In particular, CAs consist of grid cells that can take different values and are updated in discrete time steps based on transitional rules depending on the neighborhood of a cell (Packard and Wolfram 1985) (Figure 1 (a)). Then, the configuration of the grid cells or matrix matters for defining and describing different spatial and temporal situations, for example local environments related to indoor locations or large scale movements associated with transport systems (Lugo and Alatriste-Contreras 2022; Lugo and Martínez-Mekler 2022). In this case, our matrix shows a closed boundary that represents real life situations in which the crowd is limited by physical boundaries, such as walls, buildings, and natural structures. These type of boundaries are closely related to locations and areas—indoors and outdoors—of crowd events, such as mass gatherings and festivals.

**Figure 1.**
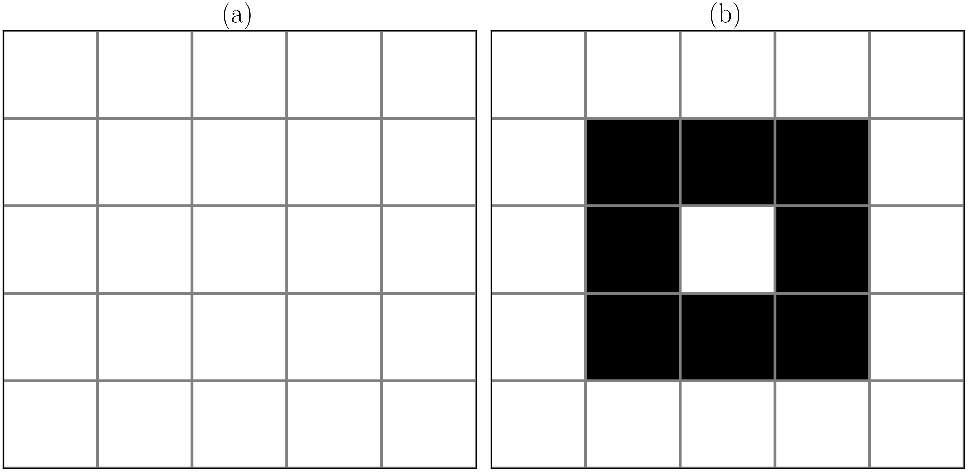
Configurations. Subfigure (a) shows a simple representation of a matrix with closed boundary. Subfigure (b) displays the shape of the neighborhood associated with the PS. The black cells represent the neighbors around a single cell.

**Figure 2.**
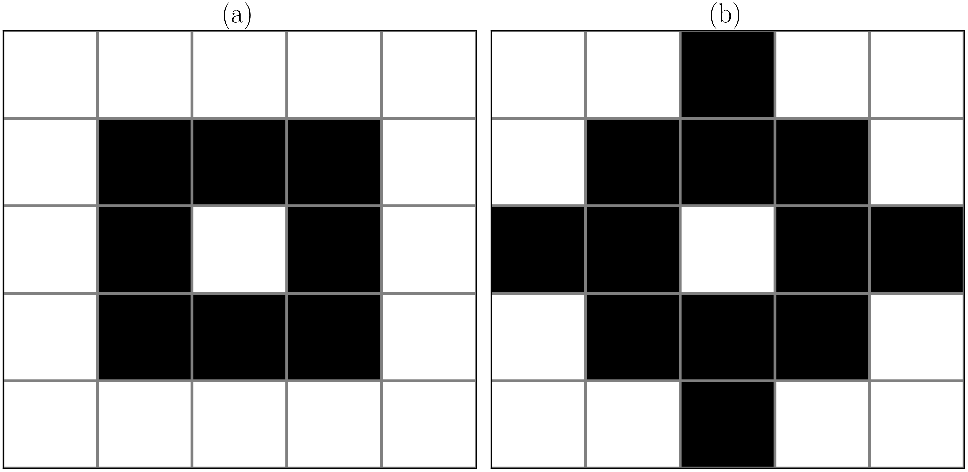
The PS and the nearest neighbors measure. Subfigure (a) displays the neighbors related to the PS. Subfigure (b) shows the neighbors associated with the computation of the nearest neighbors measure.

Next, each individual will be placed in each cell and move or stay in place based on the presence of surrounding individuals that occupied a cell and their level of tolerance (Figure 1 (b)). Based on the Shelling’s model of segregation, we shall see the emergence of group behavior associated with crushing events as a result of relative and modest decisions of individuals who act in their self-interest and group preference (Batty 2005). In this respect, we consider the concept of “phycological crowd” for describing the presence of different groups of people within a physical crowd linked by a clear social coherence (Reicher 1984; Reicher and Drury 2011). Then, in this case, we can define individuals as crowd members who are identified with each other based on their similar opinion or preference (Templeton et al. 2015). This preference can be associated with two values of occupied cells by individuals that define which individual shows the attribute of “tolerance” or “intolerance” of being surrounded by others. In addition, each individual shows the attribute of movement around its unoccupied cells. This attribute considers the possibility of evading the formation of a self-overpopulated neighborhood or staying with similar ones. Therefore, each cell shows the attribute of occupied (tolerance or intolerance) and unoccupied cell.

Let us now consider the neighborhood of an individual. In particular, we used the Moore neighborhood to describe the PS (Figure 1 (b)). Because the PS can be seen as a “circular area surrounding the person” (Welsch et al. 2019), the Moore neighborhood can be a good approximation for modeling the interpersonal distance around a person in near-body or extreme close interactions (Packard and Wolfram 1985; Hayduk 1978). Furthermore, using this type of neighborhood and the Shelling’s model of segregation about the decisions of individuals based on similar surrounding preferences (Schelling 1978), we can associate the concept of “social psychology” with the “sociocultural approach” in crowd situations (Vygotsky 1978; Reicher 1984). In particular, the behavior and thoughts of an individual to move or stay in place in a crowd can be influenced by the cultural context— family, friends, communities, and religion. Then, the socio-cultural environment of the place in which the crowd is gathering affects the instinctive reaction of people to move or stay in place. Therefore, based on these ideas, we can identify and set the number of neighbors who affects the preference of an individual of being surrounded by its similars or moving to other unoccupied cell due to its preference of avoiding near-body proximities with other individuals. These situations can describe the individual psychology behind the physical reaction of a person within a crowd.

### Rules

This subsection describes the transitional rules related to the individual behavior based on itself-interest and group behaviors. Following the notation in Batty (2005), we defined each cell in a *N* x*N* matrix, *m*_*NxN*_, as *C*_*ij*_(*t*) in which each cell, *C*, shows the location (*i, j*) at time *t*. The notation (*i, j*) is referred as a tuple of values related to the coordinates of *m*, and *t* is related to the number of iteration for updating *m*. Each *C*_*ij*_(*t*) has the attribute of occupied or unoccupied cells. Occupied cells are related to individuals who have the attribute of tolerance or intolerance as 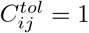 or 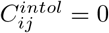 respectively. On the other hand, unoccupied cells are defined as 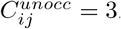. The attributes of tolerance and intolerance will not change as *t* moves forward. The change of each individual will be in its location (*i, j*) based on its neighbors. We defined a neighborhood of eight cells in which the number of neighbors related to tolerance or intolerance individuals are the following:

where *k*_*ij*_ is a cell location occupied by an individual in the neighborhood Ω, and *k*_*ij*_ shows the attribute of tolerance or intolerance.

Then, the movement of an individual is related to the number of unoccupied cells and the number of similar individuals around it. Therefore, we updated each cell as follows:

where 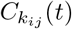 is a neighbor cell; 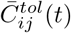 and 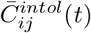 are the sum of the values of neighbors related to an individual with tolerance and intolerance attributes respectively; and *tol cells* and *untol cells* are parameters for specifying different levels of tolerance and intolerance—i.e., high and medium levels. Following the ideas of Thompson et al. (1979) and Welsch et al. (2019) that there is an immediate discomfort associated with a near-body or extreme close proximity presence, we describe equation 2 as follows. The first and second situations in equation 2 set the rules of movement and immobility of an individual based on the existence of unoccupied cell and similar individuals around it. In particular, the first and second rules describe situations in which an individual with a tolerance or intolerance attribute will move its location around its neighborhood or stay in place based on the number of available cells and the number of similar individuals around it. The third rule describes a situation in which an individual does not move due to the lack of empty space around it.

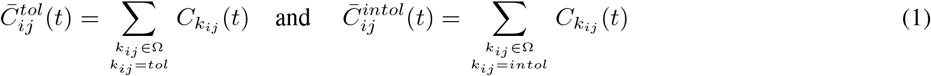

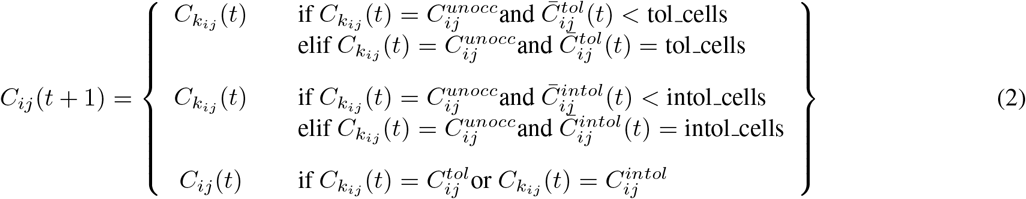

Based on the parameters *tol cells* and *untol cells*, we can define three types of criteria related to similar number of individuals around it, and each of them described levels of tolerance or intolerance. The first criterion is related to a similar level—medium level—of tolerance and intolerance among individuals (Algorithm 1). This criterion describes a situation in which the discomfort associated with the violation of the PS is the same for the two types of individuals. In particular, in equation 2, we set the parameters as follows: *tol cells* = 4 and *untol cells* = 4.

The second criterion is related to different levels of tolerance and intolerance among individuals (Algorithm 2, see the Supporting information). In particular, for tolerant individual, we specify the variable *tol cells* = 7, and for intolerant ones, we set *untol cells* = 4. This situation describes the case of high levels of tolerant individuals who will move their location or stay in place until they are almost caught by neighbors. Meanwhile, the intolerant individuals show a medium level.

The third criterion is associated with extreme levels of tolerance and intolerance among individuals (Algorithm 3, see the Supporting information). In particular, for tolerant individuals, we specify the variable *tol cells* = 7, and for intolerant ones, we set *untol cells* = 2. This situation describes the case of high levels of tolerant and intolerant individuals.

#### Algorithm 1

Rules for moving or staying in place based on similar tolerant and intolerant individuals

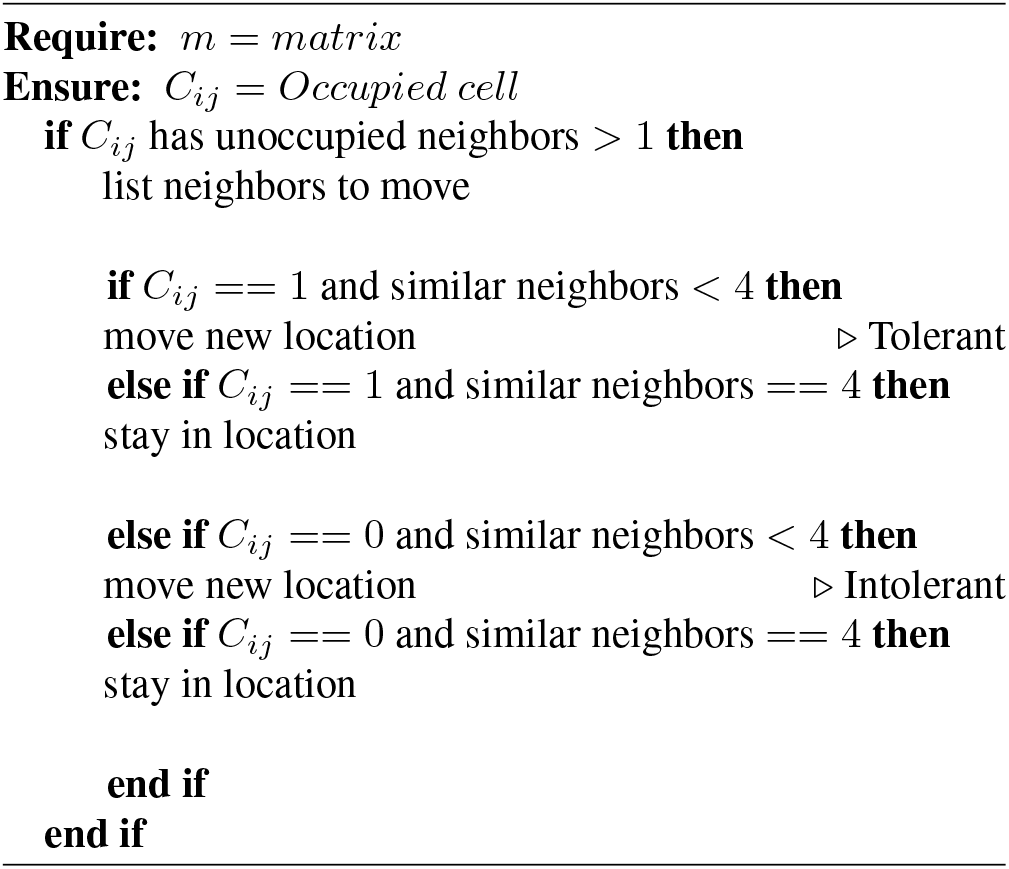

### Nearest neighbors measure and density scenarios

In line with the PS and the possible outputs associated with the rules above, we compute a nearest neighbors measure that aims to describe a local density of people around an individual. We use a neighborhood of 12 cells for counting the closest neighbors (Moussaid et al. 2011). In terms of the PS, this measure shows how well or badly the individual manages the self-perception of comfort or discomfort in a group behavior. For example, if the measure shows one or two persons around, the individual could well manage the self-perception of comfort or discomfort without threatening its security. Therefore, this measure is computed by a function named *cnt neighbors* (see the OSF project for details on the programming code: Cultural drivers for understanding crowd crushing).

The nearest neighbors measure is one key to identify a dangerous and potentially fatal situation because an individual can be jammed by its 8 or 12 nearby and occupied cells. This measure can provide critical information of possible crushing situations; no matter if your neighbors are similar or dissimilar, you are in a dangerous location position, if you are jammed by people.

Next, because we are interested in exploring different levels of crowd density, we incorporate into the initial conditions different scenarios of them. These are related to proportions of occupied or unoccupied cells by individuals in a fixed NxN matrix. As we mentioned in subsections of Configuration and Rules, we consider three types of cell attributes, 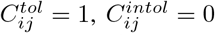, and 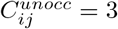, that are used for specifying those proportions. In particular, depending of the initial matrix dimensions, we can set the value of each cell randomly based on particular probabilities associated with those cell attributes. Instead of using a uniform distribution in which there are the same probabilities of selecting those attributes, we generate a non-uniform random sample based on those attributes (see the OSF project for details on the programming code: Cultural drivers for understanding crowd crushing). For example, if we generate a 10×10 matrix based on the values of 0, 1, or 2, we can define one scenario in which the occupied cells are more than the unoccupied cell as follows: (0.4, 0.4, 0.2). The first decimal is the probability associated with occupied cells with the tolerance attribute, the second decimal is the probability related to occupied cells with the intolerance attribute, and the third value is the probability associated with the presence of unoccupied cells, (*Occupied*_*tol*_,*Occupied*_*intol*_, *Unoccupied*). Such a tuple of decimals must sum 1 for purposes of normalization. Then, the tuple (0.4, 0.4, 0.2) defines a density of 80% in a total of 100 number of cells. Therefore, we aim to explore three types of density levels that covers the most generative cases, density of 40%(0.2, 0.2, 0.6), 60% (0.3, 0.3, 0.3), and 80%(0.4, 0.4, 0.2).

### Data analysis and model validation

Based on the above nearest neighbors measure and density scenarios, we apply an explorative data analysis and model validation (Sargent 1984; Leek and Peng 2015). In particular, the exploratory data analysis aims to search for trends and relationships between our nearest neighbors measure and the density escenarios for explaining the public perceptions and scientific knowledge about the crushing events. For example, how true is the public perception of an increasing concern about the dangers of overcrowding events? and how accurate is our local measure for describing harmful effects in overcrowded events? Then, to determine whether our theoretical model describes the group behavior and the presence of crushing events, we should evaluate the dynamics of the nearest neighbors based on different set of initial conditions (not only in the density scenarios, but also in parameters related to the variable levels of tolerance or intolerance in the transitional rules).

We propose a conceptual model validation in which the theory and assumptions generate consistent and justifiable results (Kerr and Goethel 2014). Then, for validating our model, we use a probabilistic framework based on a Monte Carlo approximation that looks for statistical regularities associated with our initial conditions (Table 4). The presence of such regularities can validate the use of our initial conditions because the model outputs may agree with the observed data of public perception of crushing events. Therefore, our model validation process shall support our conclusions.

In particular, we use a sensitivity analysis that determines the relative influence of the initial conditions and the parameters on our model output. We are interested in exploring which combination of those parameters are related to the current public perceptions and scientific knowledge of crushing events. To do this, we shall follow the procedure outlined below (Figure 3).

**Figure 3.**
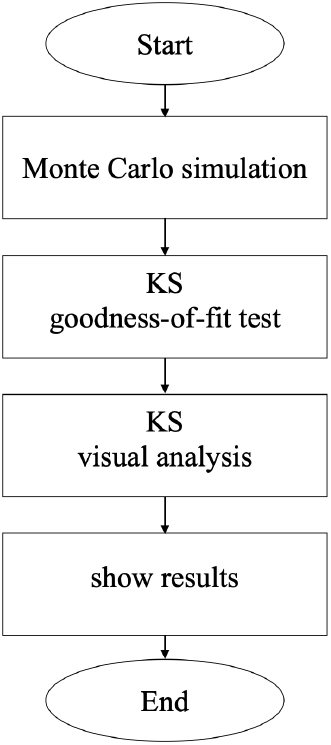
Flowchart of the model validation.

**Figure 4.**
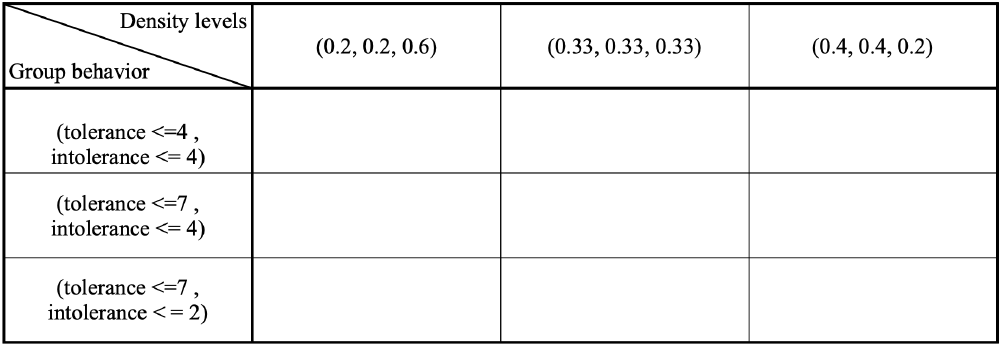
Initial conditions by density and group behavior. Based on the density levels and the group behavior, this table shows the main scenarios for exploring in our model. For simplicity in the data analysis and model validation, we organize the results based on this table.

**Figure 5.**
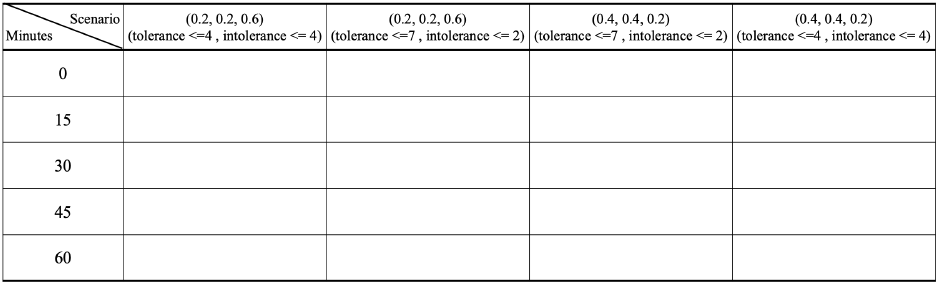
Initial conditions by minutes and scenarios. Each column shows an scenario related to particular values of density levels and the group behaviors. Each row displays the a particular time in minutes. For simplicity in the data analysis and model validation, we organize the results based on this table.

**Figure 6.**
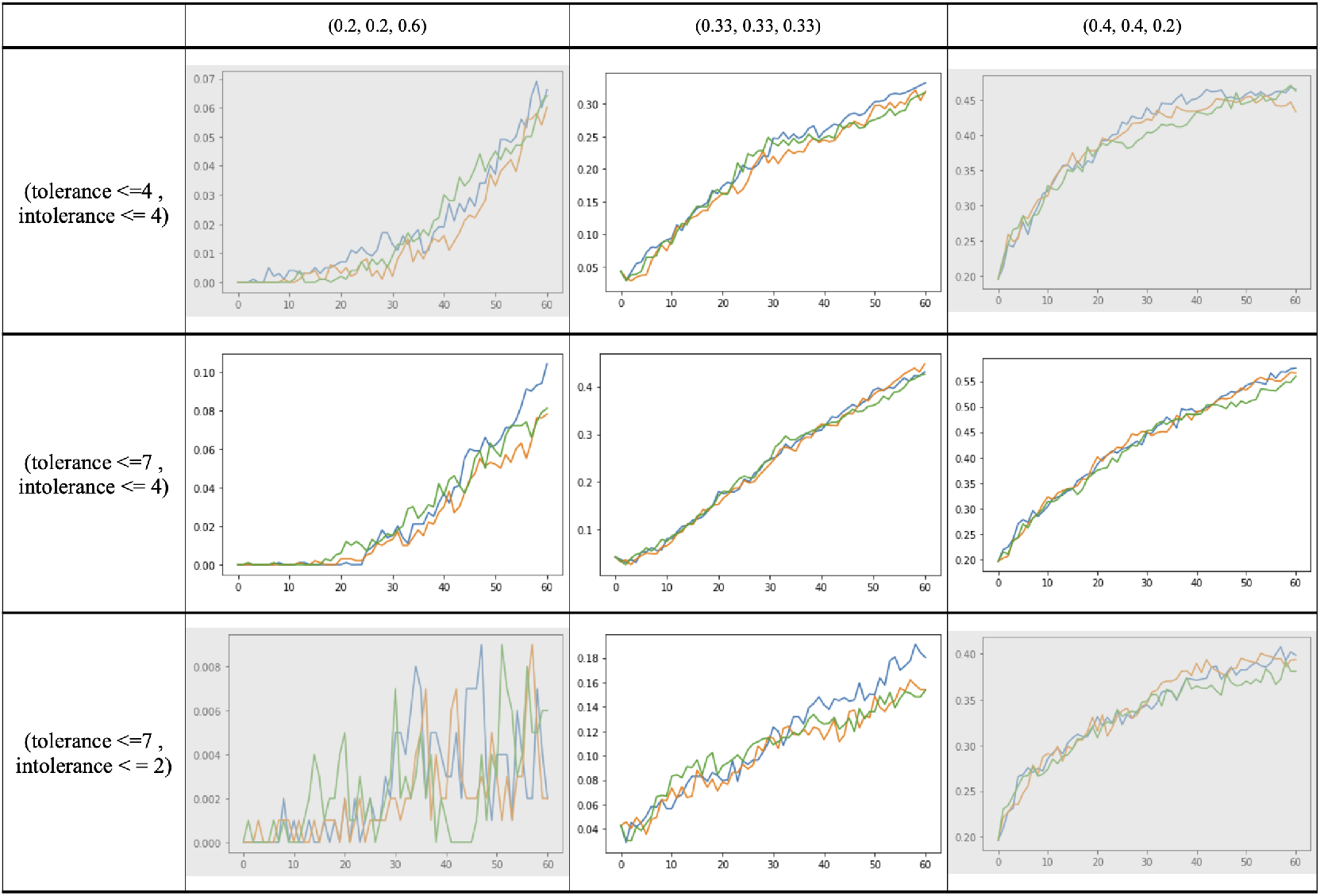
Sensitivity analysis. The random seeds were set to obtain the same initial values of the matrix. For exemplifying the results of a total of 10, 000 realizations, each subfigure display three realizations per case. Subfigures in rows show variations in the group behaviors (tolerance and intolerance parameters), and columns show variations in the proportion of occupied and unoccupied cell parameters (density levels). The *x* axis in each subplot shows the number of iterations, and the *y* axis shows the values of the nearest neighbors measure based on cells of 12 surrounded neighbors. The shaded boxes represent the cases that we use for an in-depth analysis in Fig 9 and Fig 13

The first process is to apply a Monte Carlo simulation. It tests how probable is the presence of crushing events in certain density situations. We set our number of trials in 10, 000. Then, for each realization related to density levels and different parameters associated with group behaviors, we present a best-fit analysis that aims to identify the statistical distribution that best describes our generated data. In particular, this second process uses the Kolmogorov-Smirnov (KS) goodness-of-fit test for identifying the statistical distributions that best describe the data (Massey 2020). This test compares each realization of data with a set of theoretical distributions. In this respect, we followed the procedure in Lugo and Martínez-Mekler (2022) and Lugo and Alatriste-Contreras (2022) who use the normal, log-normal, exponential, Pareto, Gilbrat, power law, and exponentiated Weibull as the main theoretical distributions. Then, we review the KS test statistics, and their visual representations, which are based on the empirical and theoretical cumulative density functions of a sample. Finally, the fourth process, we report our results.

### Initial conditions

Based on the above processes, we explore different initial conditions related to density levels and parameters of tolerance or intolerance in the transitional rules. For simplicity, the total number of cells is set by a matrix with fixed dimension equal to (50, 50), that is 2500 cells. Recall that this matrix shows a closed boundary that represents physical obstacles for free movements. It is important to note that the scale of the analysis can represent accurately increasing scales. Therefore, there is a consistency between our theoretical assumptions, the generated synthetic data, and the public perceptions and scientific knowledge about the crushing events.

As we mentioned above, we define the following set of tuples related to different scenarios based on the proportions of tolerant, intolerant, and unoccupied cells respectively: (0.33, 0.33, 0.34), (0.2, 0.2, 0.6), and (0.4, 0.4, 0.2). The first scenario is related to similar proportions between occupied and unoccupied cells; the second is associated with decreasing and similar proportions of occupied cells, but an increasing proportion of unoccupied cells; and the third is related to increasing and similar occupied cells, but a decreasing proportion of unoccupied cells (Table 4).

In addition to these density levels and based on the three criteria mentioned in the Rules subsection, we are interested in the nature of relationships between individuals in a group. Then, we explore three sets of variations in parameters related to tolerant and intolerant individuals (Table 4, 5). The first is related to a situation in which the discomfort or comfort associated with the violation of the PS is the same for tolerant or intolerant individuals. The second is a circumstance where tolerant individuals show a high level of confort associated with the violation of the PS. The third situation is related to a presence of high levels of comfort and discomfort associated with tolerant and intolerant individuals respectively.

Finally, the number of iterations in each realization is fixed to a total of 60. We set this value as a reference to time, in minutes (Table 5). In particular, we assumed that the presence of crushing events can happen within a range of 60 minutes, in a window of 15 min. (Moussaid et al. 2011; The Times of Israel 2021; BBC 2022).

## Results

In this section, we present our findings obtained from the explorative data analysis. In particular, we display graphical and statistical results based on our model validation. We use the sensitivity analysis for exploring the dynamics of the nearest neighbors measure based on different sets of initial conditions and parameters. These sets aim to identify the current public perceptions and scientific knowledge about the importance of being tolerant or intolerant to similar ones and the consequence of the interpersonal distance in overcrowding events. We display the results based on Tables 4 and 5 for drawing attention to the most representative of cases.

Fig 7 shows the general results of the sensibility analysis that point out the importance of density situations over the group behaviors. There is evidence that variations in density levels affect considerably the dynamics that generate crushing events. For example, if we select the first row of the table, which is characterized by similar behaviors of individuals, we can see that increasing the proportion of unoccupied cells affects the nearest neighbors measure showing overcrowding values. That is, the first entry in this row shows the lowest values of the nearest neighbors measure. This indicates that if we control the number of people in a crowd by giving more space or areas in which people can move without jamming, the probability of generating crushing is lower (see Fig 9).

**Figure 7.** Example header text.

**Figure 8.**
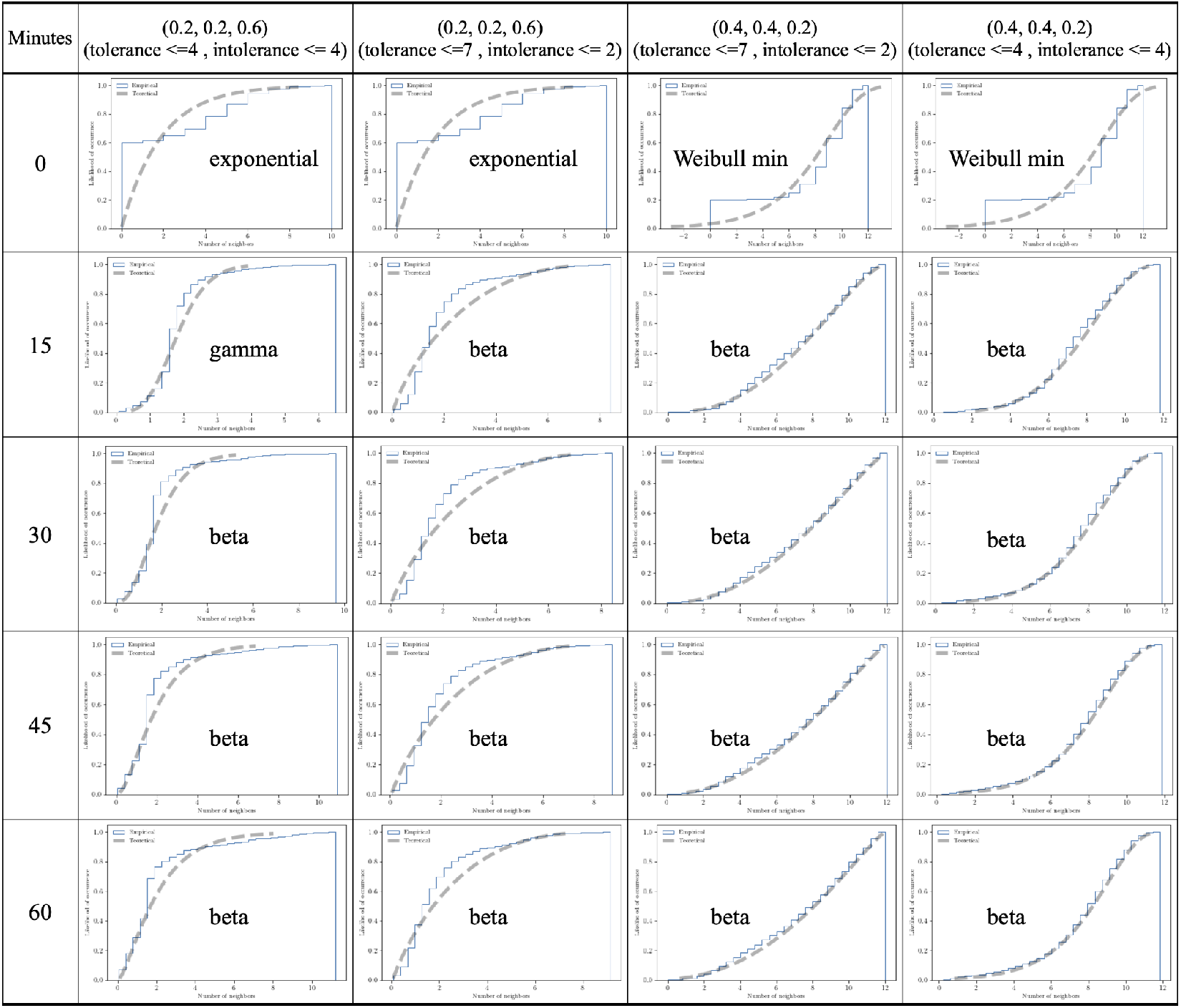
Best-fit analysis of fourth selected cases of the sensitivity analysis at particular given time step. Subfigures are the visual results of the KS test (see Table 1, supplementary material, for seeing the probability density functions, PDF). These results are based on a Monte Carlo simulation of a total of 10, 000 realizations per case. Subfigures in rows show particular time in minutes, and columns shows the selected cases of the sensibility analysis. The *x* axis in each subplot shows the value of the nearest neighbors measure, and the *y* axis shows the likelihood of occurrence of particular values of the nearest neighbors measure.

**Figure 9.** Example header text.

**Figure 10.**
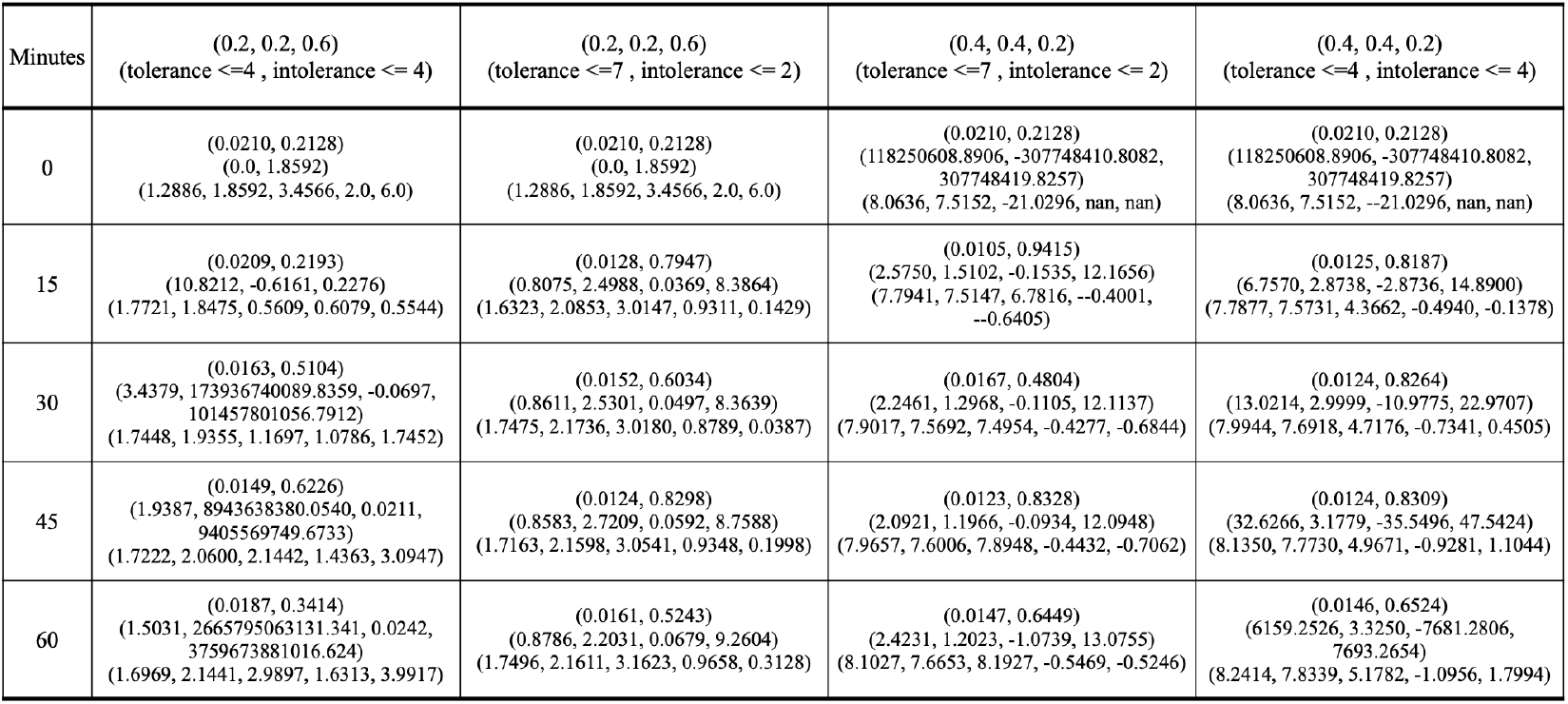
KS test, estimated parameters, and first moments of the selected cases. KS test: (stat, p-value), estimated parameters based on the KS test: (parameter1, loc, scale), first moments: (median, mean, variance, skewness, kurtosis). These results are based on a Monte Carlo simulation of a total of 10, 000 realizations per case.

**Figure 11.** Example header text.

**Figure 12.**
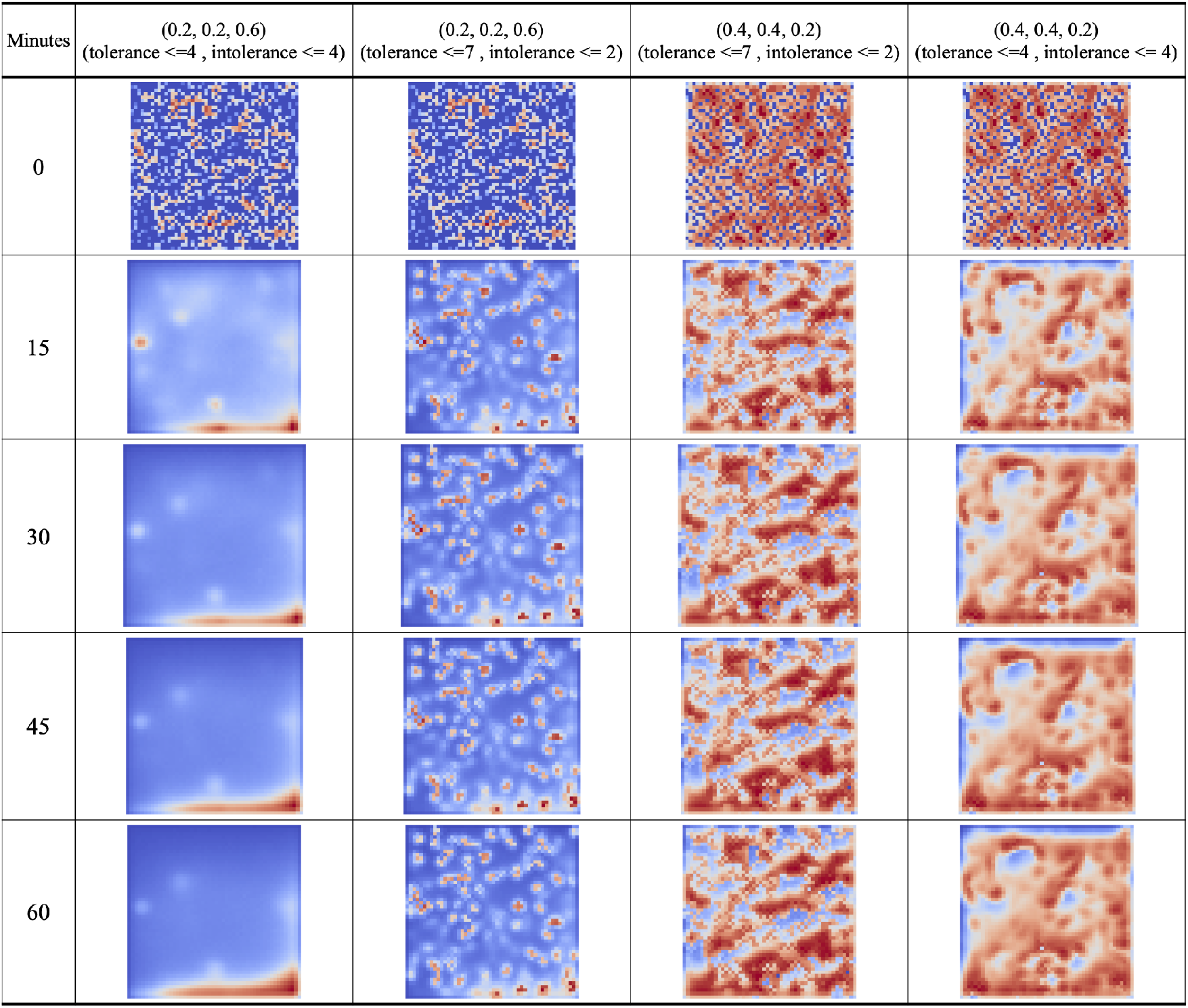
Nearest neighbors measure of selected cases of the sensitivity analysis at particular time. We use the “coolwarm” palette to display the result. From dark blues to bright reds, we can see the range of number of neighbors, from zero to 12 neighbors. These results are based on a Monte Carlo simulation of a total of 10, 000 realizations per case. Subfigures in rows show particular time in minutes, and columns shows selected cases of the sensibility analysis.

Now, if we see the third row, which is characterized by dissimilar behaviors of individuals—i.e., extremist positions of tolerant or intolerant individuals—we can see a sharp drop in the proportion of cells with high values of the nearest neighbors measure. This result is particularly important because it suggests that even if there is an extreme preference of being surrounded by others, the drastic position of being intolerant produces lower values of the nearest neighbors measure. That is, the crushing events are less probable (see Fig 9).

The columns 2 and 3 in Fig 7 display the presence of unstoppable crushing events. All of these cases show a marked increase in values of the nearest neighbors measure. Even if there is a similar density between occupied and unoccupied cells, as well as the presence of extremist preferences of being tolerant or intolerant, the values of the nearest neighbors measure rapidly and dangerously increase. These results suggest that crushing events are more probable than we have thought at the beginning (see Fig 9).

With respect to our fourth particular case, we can see the behavior of the nearest neighbors measure in different time periods. Fig 9, 13, and Table 11 display the KS test per time period related to the nearest neighbors measure. Fig 9 reveals that there is a clear difference between the best-fit statistical distributions of the initial conditions and the rest of the periods. It suggests a differentiated behavior of individuals that produce a marked transition between statistical distributions. Furthermore, this Fig 9 shows that there has been a marked presence of the beta distribution in most of the periods of time. The preponderance of these results suggests the presence of highly skewed statistical distributions behind this type of collective behavior.

**Figure 13.** Example header text.

Finally, we emphasize that the case with the lowest probability of generating crushes is the one where there are dissimilar behaviors of individuals and a lower density of unoccupied cells. This result is clearly displayed in column two of Fig 9 and 13. This particular case shows a small number of dispersed clusters of individuals in all periods of time. It suggests that this particular situation delays the process of crushing events.

Therefore, it seems that the density related to the proportion of unoccupied cells matters the most. An increased proportion of unoccupied cells or lower density is more preferable than an increase proportion of individuals with extreme preferences among them.

## Discussion

As previously stated in the Introduction section, our main questions aimed to identify situations in which the crushing is delayed, to show how probable are the presence of crushing events, and to differentiate the group behavior of tolerant and intolerant individuals surrounded by similar ones. Based on our findings, we reach the following interpretations.

Depending more on the initial density than the group behaviors of extremist preferences of individuals of being surrounded by similars, the crushing events are unstoppable and highly probable in dense crowd situations. The number of unoccupied cells can diminish the presence of crushing events, meanwhile the group behavior of tolerant and intolerant individuals control how fast the crushing events can develop. Contrary to expectations, the only presence of extreme preferences of being tolerant or intolerant does not ensure an effective control of crushing incidents in dense crowd situations. Therefore, our hypothesis is partially true, this would imply that differences in individual preferences for being surrounded by people in near-body proximities may explain the generation speed of crushes, but unfortunately they cannot prevent or stop them.

The fact that the beta appeared as best fit in most escenarios may represent a statistical regularity that provides a mathematical model based on frequent results. This model could be used for predicting possible outcomes and computing occurrence probabilities.

These findings have implications for understanding the dynamics behind dense crowd events. In particular, no matter the type and scale of social gatherings, it is indisputable the presence of uncontrollable factors that can cause severe personal injuries. Social gatherings, indoor or outdoors, can represent the perfect situation for a fatal crushing.

Therefore, we suggest the following actionable guidelines for crowd control professionals and nonprofessionals:

i. To organize an event in which the crowd density is related to 40% of individuals and 60% of free space.
ii. To organize an event in flat areas with a smaller number of physical obstacles.
iii. To organize an event with a higher number of emergency exits and scape routes.
iv. To inform assistants the importance of the crowd density and the risk of being crushed.
v. To inform assistants to lookout every 15 minutes for any sing of chaotic behavior around them.

With specific reference to the computational efficiency or scalability of our theoretical model, we can see stability in the processing time of each execution per scenario. In particular, if the size of the array increases, the processing time increases too. Even though these variations of the initial matrices, from small to large scales, the dynamics of the system remain similar. Therefore, our model shows consistency between the public perception and our simple modeling of crushing disasters. Our model is useful for the purpose of describing the effects of the PS in dense crowd scenarios.

Despite these promising results, questions remain. In particular, what are the effective practices for communicating the risks of social gatherings? How the current technology of mobil applications can prevent possible crushing in social gatherings? Future work is needed to generate an application for predicting fatal situations. We are considering to explore the generation of a warning signal in which every mobil device can be used to prevent fatal crushing. Similar to the warning signal of earthquakes, it is possible to generate a mobile application that indicates a warning signal of probable crushing events in real time based on the nearest neighbors measure and density.

In addition, it is important to generate simple and generative models that consider the presence of external factors, such as weather, time of day, and event time. These factors can explore the effects of the generation speed of crushes.

## Conclusion

Different types and scales of social gatherings in progress might show high probability of occurrence of crushing events. The self-awareness of your personal space plays a vital role for preventing risky situations and reacting during overcrowding. An initial situation in which there is more space than attendees in a social gathering is preferable, and it could reduce the risk of crushing events. When this condition is not feasible, there are high probabilities of harmful incidents and fatal crushing events. Therefore, every coordination committee of those type of social events should communicate the risk of being hurt due to the lack of enough room for moving freely.

## Data availability

The research data and the programming code is available in the following open data repository: Open Science Framework, project named Cultural drivers for understanding crowd crushing.

## Appendix

### Algorithm 2

Rules for moving or staying in place based on high tolerant and medium intolerant cells

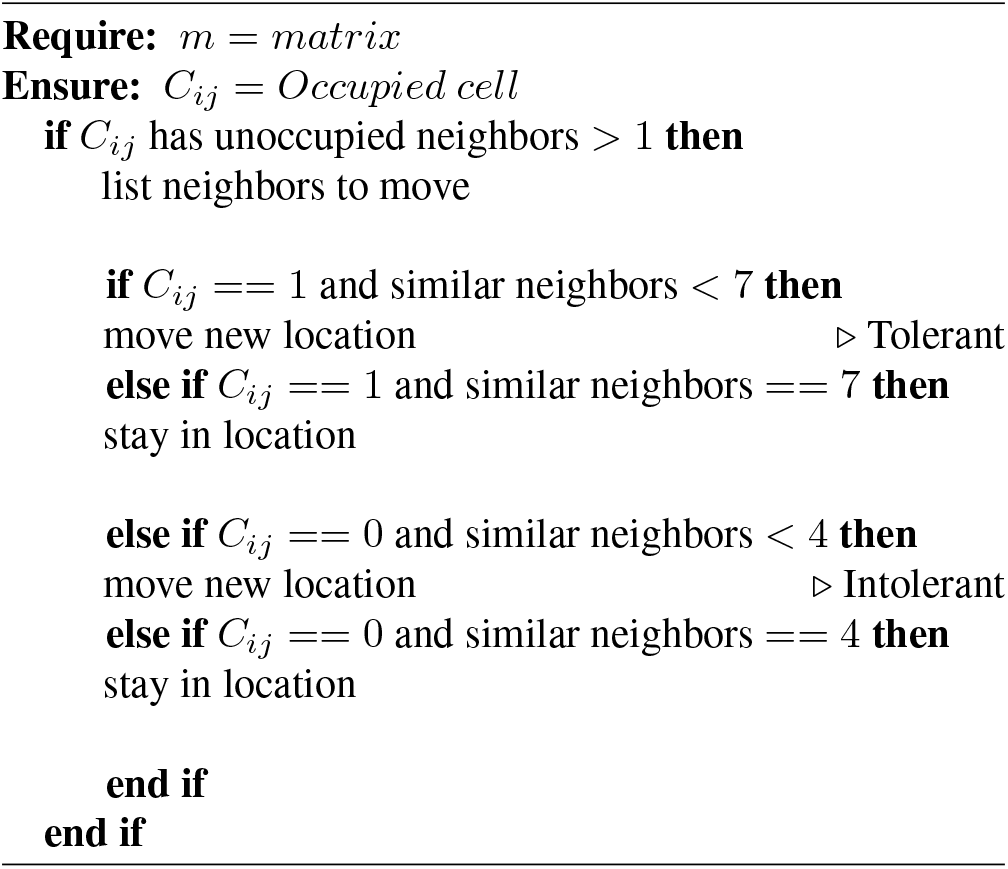

**Table 1.** Statistical distributions of the KS test and their probability density functions (PDF). Name of the statistical distributions and their PDF used for the KS test.

| Name | PDF |
| --- | --- |
| exponential | $f(x) = \exp(-x),$<br>for $x \geq 0$ |
| gamma | $f(x, a) = \frac{x^{a-1}e^{-x}}{\Gamma(a)},$<br>for $x \geq 0$ and $a > 0$ |
| beta | $f(x, a, b) = \frac{\Gamma(a+b)x^{a-1}(1-x)^{b-1}}{\Gamma(a)\Gamma(b)},$<br>for $0 \leq x \leq 1, a > 0, b > 0$ |
| Weibull minimum | $f(x, c) = cx^{c-1}\exp(-x^c),$<br>for $x > 0, c > 0$ |

### Algorithm 3

Rules for moving or staying in place based on high tolerant and high intolerant cells

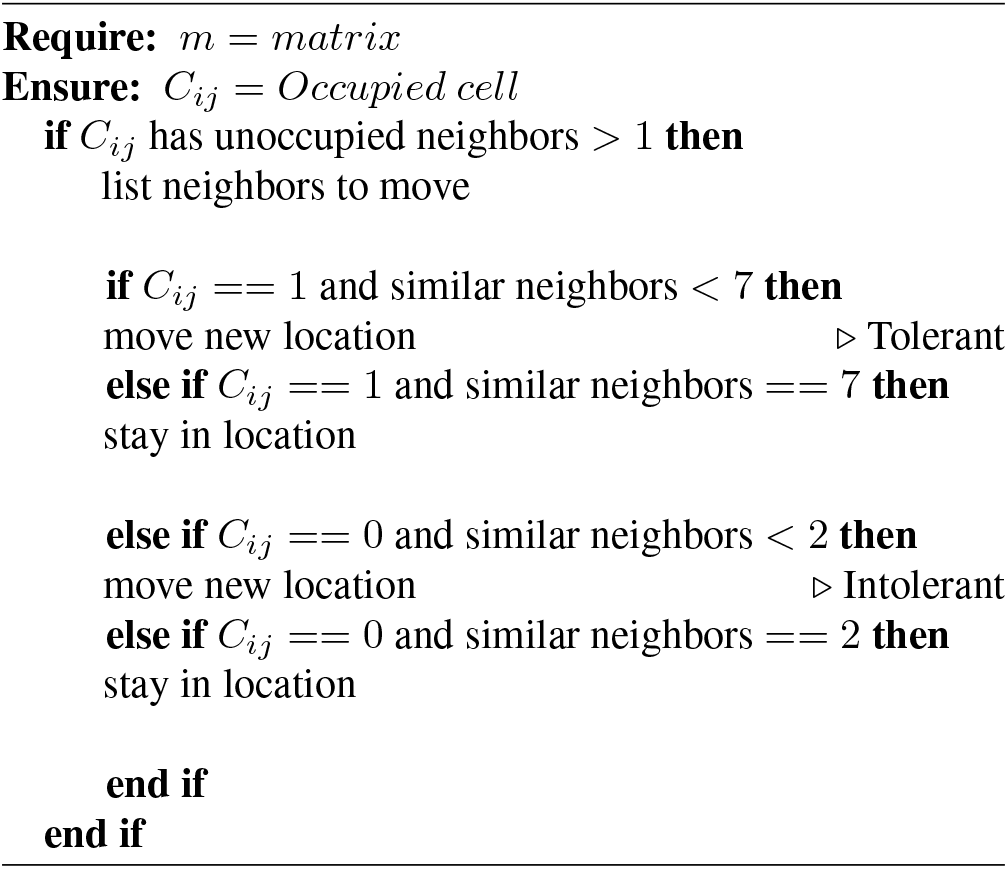

